# Target cell tension regulates macrophage trogocytosis

**DOI:** 10.1101/2024.12.02.626490

**Authors:** Caitlin E. Cornell, Aymeric Chorlay, Deepak Krishnamurthy, Nicholas R. Martin, Lucia Baldauf, Daniel A. Fletcher

## Abstract

Macrophages are known to engulf small membrane fragments, or trogocytose, target cells and pathogens, rather than fully phagocytose them. However, little is known about what causes macrophages to choose trogocytosis versus phagocytosis. Here, we report that cortical tension of target cells is a key regulator of macrophage trogocytosis. At low tension, macrophages will preferentially trogocytose antibody-opsonized cells, while at high tension they tend towards phagocytosis. Using model vesicles, we demonstrate that macrophages will rapidly switch from trogocytosis to phagocytosis when membrane tension is increased. Stiffening the cortex of target cells also biases macrophages to phagocytose them, a trend that can be countered by increasing antibody surface density and is captured in a mechanical model of trogocytosis. This work suggests that a distinct molecular pathway for trogocytosis is not required to explain differences in trogocytosis among target cell types and points to a mechanism for target cells to modulate trogocytosis.

## Introduction

Phagocytosis by macrophages, first observed by Metchnikoff more than 140 years ago^1^, plays a critical role in the immune system. Upon binding and recognition of unhealthy cells or foreign particles, macrophages can fully engulf and destroy the targets, contributing to the clearance of bacterial infections, cell corpses, and, more recently, immunotherapy-targeted cancer cells^2,4^. One of the best characterized mechanisms for targeting a particle for engulfment by macrophages is antibody dependent cellular phagocytosis (ADCP). Once labeled with antibodies, pathogens and other cells in the body are recognized by Fc-receptors (FcR) on the macrophage surface, triggering a signaling cascade that results in growth and closure of an actin-driven phagocytic cup around the target, typically followed by destruction after internalization.

More recently, macrophages have been observed to sometimes simply ‘nibble’ portions off of the target cell membrane rather than fully engulf the target^5,6^. This peculiar process, termed ‘trogocytosis’ (*Trogo*-; Greek for ‘to nibble’), is emerging as an important effector function on its own that can both enhance and undermine immune cell function^7^.

For example, macrophages have been observed to nibble hematopoietic stem cells, marking them for retention in the bone marrow^8^. Similarly, tissue resident macrophages of the brain, microglia, selectively nibble synapses of neurons to prune connections in the developing mouse brain^9^. However, trogocytosis is sometimes an undesirable outcome, especially in the innate immune response to cancer immunotherapies. Instead of phagocytosing cancer cells target with therapeutic antibodies, macrophages will remove the antibodies from the cell surface via trogocytosis, leaving the cancer cells alive^5,10,11^.

Beyond the mammalian immune system, trogocytosis has been observed in cell types as diverse as the pathogenic amoeba *Entamoeba histolytica* and the endodermal cells of *C. elegans*. The role of trogocytosis in these organisms is similarly diverse. For pathogenic *Entamoeba*, amoeba will first kill a cell via trogocytosis and then fully phagocytose their deceased target^12^. Interestingly, the amoeba will subsequently display proteins from the target cell, a process known as ‘cross-dressing’, cloaking themselves from host immune attack^13^. Trogocytosis plays a gentler homeostatic role in *C. elegans*; lobes rich in mitochondria on primordial germ cells are pruned by endodermal cells in close proximity, potentially reducing the risk of reactive oxygen species in the germline^14^.

Despite the importance of trogocytosis, the signals on the target cell that drive trogocytosis over phagocytosis or other cell-cell interactions are poorly understood. It is unlikely that this signal is a conserved cell surface molecule, antibody, or soluble cytokine, as diverse target cells in a wide range of physiological systems appear to be able to promote trogocytosis. It is also unlikely to be a purely stochastic phenomenon, as not all cell-cell interactions, for example between macrophages and apoptotic cells, lead to trogocytosis as well as phagocytosis. For the primordial germ cells of *C. elegans*, cellular markers of distress (*e.g.* reactive oxygen species) seem to mark cells for trogocytosis^14^; however, this has not been shown to be the case for microglia or pathogenic amoeba.

Here we find that the physical properties of the macrophage-target interface, rather than specific molecular components of the target surface, drive the macrophage decision to trogocytose rather than phagocytose. We quantitatively evaluate the trogocytic and phagocytic efficiency of macrophages interacting with antibody-labeled cells and membrane-only cell mimics using a combination of light microscopy, micropipette aspiration, and flow cytometry. Using giant unilamellar vesicles (GUVs), we confirm that membrane tension is a sufficient signal to bias trogocytosis over phagocytosis by varying tension in the absence of surface molecules other than antibodies. We further demonstrate that trogocytic efficiency of antibody-labeled cells depends on target cortical tension and the density of antibodies coating the target cell surface. Finally, we propose a mechanical model of macrophage trogocytosis that accounts for the observed dependence on target surface tension and antibody density.

This work shows that a distinct molecular pathway is not required to explain why some target cells are trogocytosed rather than phagocytosed. Instead, the macrophage’s ability to deform the target cell surface after binding is a sufficient signal to promote trogocytosis, suggesting that phagocytosis may be thought of as frustrated trogocytosis.

The increase in trogocytosis with decreased cell cortical stiffness raises the possibility that tumor cells and other phagocytic targets could modulate their physical properties to evade phagocytosis.

## Materials

### Lipids

Phosphocholine (PC) lipids (Avanti Polar Lipids) were used as purchased without further purification. Lipid stock solutions in chloroform contained a quaternary mixture of 97.5 mol% POPC, 1 mol% biotin-PE, 1 mol% PEG2K DSPE, and 0.5 mol% lissamine rhodamine PE. GUVs are diluted in an ionic solution of PBS and all lipids in our mixtures are zwitterionic. We added PEG2K DSPE to block GUVs from aggregating in the charge- screened PBS solution.

### Antibodies

Antibodies used to opsonize GUVs and cultured cells were purchased from Santa Cruz Biotechnology and used without further labeling or purification. Biotin and CD47 were bound by, respectively, AlexaFluor647-labeled anti-biotin mouse IgG (clone BK-1/39, Santa Cruz Biotechnologies) and AlexaFluor647-labeled anti-CD47 mouse IgG (B6H12, Santa Cruz Biotechnologies). The Fc-receptors on RAW cells were bound by anti- CD16/32 mouse IgG (Biolegend).

### Methods

### RAW 264.7, J774A.1, Jurkat T, Raji B, and HL60 cell culture

RAW 264.7 murine male macrophage-like cell line, J774A.1 murine female macrophage-like cell line, Jurkat T human cell line, Raji B human B cell line, and HL60 human promyeloblast cell line was obtained from the UC Berkeley Cell Culture Facility. RAW 264.7, J774A.1, Jurkat T, and HL60 cells were cultured in RPMI 1640 media (Corning) supplemented with 10% heat-inactivated fetal bovine serum (HI-FBS, Thermo Fisher Scientific) and 1% Pen-Strep (Thermo Fisher Scientific). Raji B cells were cultured in RPMI 1640 media supplemented with 10% HI-FBS, 1% Sodium Pyruvate (Thermo Fisher Scientific), and 1% Pen-Strep. RAW 264.7 and J774A.1 cells were cultured in non-tissue culture-treated 10 cm dishes (VWR); all other cell lines were cultured in tissue culture-treated 10 cm dishes (Corning) at 37°C, 5% CO2. Pre-stimulated RAW 264.7 cells were incubated with 100 ng/mL lipopolysaccharide (LPS; Sigma) for 24 hours prior to co-culture with target cells.

### Bone marrow derived macrophages

Bone marrow derived macrophages (BMDMs) from male C57BL/6 (B6) mice were a kind gift from the Portnoy Lab (UC Berkeley). BMDMs were grown in RPMI 1640 media supplemented with 10% HI-FBS and 1% Pen-Strep at 37°C, 5% CO2. BMDMs were used in experiments within 24 hours of thawing.

### GUV electroformation

Solutions containing 0.25 mg total lipids were spread evenly on slides coated with indium tin oxide (70-100 Ω/sq; Sigma Aldrich). The slides were placed under vacuum for >30 min to allow for complete evaporation of chloroform. A capacitor was created by sandwiching a 0.3-mm rubber septum between two lipid-coated slides. The gap was filled with ∼200 μL of 285 mM sucrose (osmotically matched to PBS using an osmometer (Precision Systems)). GUVs 10 to 100 μm in diameter were electroformed^15^ by application of an AC voltage of 1.5 V at 10 Hz across the capacitor for 1 h at 55°C.

### Size-separation of GUVs

Sucrose solutions containing electroformed GUVs were diluted in 1 mL of 285 mM glucose and injected into a home-built differential density column, lovingly named the SeparatorMax 3000 (Fig. S1). Specifically, 10 ports of an inner diameter 2 cm, spaced 2.5 cm apart, were drilled into a cast acrylic tube of 33.6 cm. This tube was connected to two different baths containing a low-density glucose solution (285 mM) and a high-density glucose solution (300 mM) in series. Solutions were loaded into the column via a peristaltic pump with vigorous hand mixing. The resulting solution column is a linear gradient of glucose density from 300 mM glucose (bottom) to 285 mM glucose (top). Vesicles were allowed to sediment overnight (approx. 12 hours) and harvested from each of the 10 ports of the column. We kept the GUVs from ports 5-8 (labeling from the top to bottom), and they had diameters from 5-20 μm. We further purified GUVs by spinning the solution at 300 x g for 10 minutes and taking the bottom 100 μL of solution. This solution was then diluted in PBS containing 4 μM AlexaFluor647-labeled anti-biotin mouse IgG (clone BK-1/39, Santa Cruz Biotechnologies). GUVs used in flow cytometer-based phagocytosis/trogocytosis assays were loaded with 1 μM soluble rhodamine dye dissolved in 285 mM sucrose. GUVs were purified from excess dye in solution by running them through the SeparatorMax 3000.

### Imaging techniques

All live cells were maintained at 37°C, 5% CO2 with a stage top incubator (Okolab) during imaging. For confocal microscopy, cells were imaged with a spinning disk confocal microscope (Eclipse Ti, Nikon) with a spinning disk (Yokogawa CSU-X, Andor), CMOS camera (Zyla, Andor), and a 60x objective (Apo TIRF, 1.49NA, oil, Nikon). The spinning disk confocal microscope was controlled with Nikon Elements (Nikon). Images were analyzed and prepared using FIJI (imagej.net/software/fiji)^16^.

### Phagocytosis/trogocytosis assays

*Phagocytosis/trogocytosis of labeled cultured cells* 100,000 macrophages were seeded in wells of a tissue-culture flat-bottom 96-well plate (Falcon) in 100 μL of RPMI 1640 medium. Post-seeding, cells were incubated at 37°C, 5% CO2 for 3-4 hours prior to target addition. 1 μM CellTracker Green CMFDA was added to stain the cytoplasm of macrophages. Cells were washed 2x with media to remove excess dye.150,000 target cells (e.g. HL60s, Jurkat T, or Raji B) were diluted in 150 μL of 1640 RPMI media containing the appropriate amount of IgG (either anti-CD47 or anti-biotin), 2 μM pHrodo succinimidyl ester (Thermo Fisher Scientific), and the appropriate amount of glutaraldehyde for drug experiments. Cells were labeled for 30 minutes at 37°C and subsequently washed 3x in media. Cells labeled with anti-biotin were first treated with surface biotinylation reagent EZ-link NHS biotin (Thermo Fisher Scientific) for 30 minutes, followed by washing 3x in 100 mM glycine in PBS pH 8.0 to quench the biotinylation reaction. After washing, 100,000 target cells were added to macrophage-seeded wells and co-incubated for 1 hour.

After incubation, wells were scraped with mini cell scrapers (Biotium) and gently pipette mixed 2-3x. Cells were immediately analyzed using an Attune NxT CytPix flow cytometer (Thermo Fisher Scientific). Cells were injected into the flow cytometer at a rate of 200 μL/min and cells were gated according to the following protocol:

A control sample of CellTracker Green CMFDA labeled macrophages were run each experiment to determine the 488-laser intensity threshold for events positive for CellTracker Green CMFDA. This sample was also used to determine background intensity levels of AlexaFluor647 (647-laser) and rhodamine (568-laser). Thresholds to establish samples positive for AlexaFluor647 and/or rhodamine were determined by running target cells labeled with AlexaFluor647 anti-CD47 IgG and pHrodo.

Events were determined positive for phagocytosis if they were positive for CellTracker Green CMFDA, pHrodo, and AlexaFluor647. Events were determined positive for trogocytosis if they were positive only for CellTracker Green CMFDA and AlexaFluor647, but not pHrodo. The trogocytic and phagocytic efficiency were computed by taking phagocytic and trogocytic events, respectively, and dividing that number by the total number of CellTracker Green CMFDA positive events.

Surface density of antibody on target cells was measured by comparing cells to calibrated beads with known numbers of AlexaFluor647 fluorophores (Quantum MESF Kits, Bangs Laboratories). Labeled antibodies from Santa Cruz Biotechnologies have 5-7 fluorophores per IgG, as per the manufacturer, so all surface densities calculated in this work considers the average diameter of the cell type (calculated from confocal micrographs) and the average fluorophore per IgG.

*Phagocytosis/trogocytosis of GUVs* 100 μL of GUVs (∼1 million GUVs counted with an impedance-based cell counter (Scepter, SigmaAldrich)) were prepared with 4 μM AlexaFluor647 anti-biotin IgG in PBS and allowed to incubate with gentle rotation for >10 minutes. After washing, GUVs were added to macrophage-seeded wells as described above. Macrophages were incubated with GUVs for 30 minutes. After incubation, wells were scraped and immediately analyzed on the flow cytometer as described above. Thresholds to establish samples positive for AlexaFluor647 and/or rhodamine were determined by running GUVs labeled with AlexaFluor647 anti-biotin IgG and containing soluble rhodamine.

### Micropipette aspiration to measure target cell cortical tension and GUV membrane tension

Micropipettes were made from capillaries (1.0 mm OD, 0.58 mm ID, 100 mm length, borosilicate glass G100-4, Harvard Apparatus) drawn out with a filament pipette puller (Sutter Instruments, Novato CA). Pipette tips were forged with an adapted microforge (MicroData Instruments, USA) to obtain a smooth opening of ∼5 μm in diameter. Pipettes were subsequently filled with 0.45 μm filtered PBS using a home-built syringe-pulling device, named The Siphonator 3000 for >30 minutes. Pipettes were treated for 15 minutes in a 0.45 μm filtered 10% bovine serum albumin (BSA) solution prior to each experiment to passivate the pipette surface and prevent unwanted adhesion of GUVs and target cells to the pipette glass.

The treated pipettes were inserted into a TransferMan 4r micromanipulator (Eppendorf) mounted on a confocal microscope (Eclipse Ti2) to facilitate the manipulation of target cells and/or GUVs for surface tension measurements. To provide controlled suction, the pipettes were connected to a syringe pump (CellTram, Eppendorf) and a pressure sensor (DP103, Validyne Engineering), which measured the suction pressure applied to the pipette.

To measure surface tension, the pipette was aligned to the target interface and increasing suction pressure was applied until a membrane tube with a length equal to the pipette radius was pulled into the pipette. Using Laplace’s law, the measured suction pressure (ΔPsuction) was used to measure the interface tension (γ):

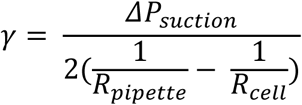

where *ΔPsuction, Rpipette*, and *Rcell* are the suction pressure, the pipette radius, and the cell radius. The interface tensions of GUVs and target cells were modulated by adjusting the suction pressure applied through the same pipettes, thereby varying the interfacial tension.

## Results

### Macrophages phagocytose and trogocytose cultured cells

To study trogocytosis by macrophages, we developed a fluorescence-based assay that uses the key features of trogocytosis (surface engulfment) and phagocytosis (surface and volume engulfment) to separate the two processes. For a volume marker, we labeled the cytosol of target cells with the pH-sensitive rhodamine derivative pHrodo, which selectively fluoresces in low pH environments (*e.g.* the phagolysosome). For a surface marker, we used a fluorescent antibody, AlexaFluor647 anti-CD47 IgG, that also activates macrophages through engagement with their Fcγ receptor (FcR). CD47 is a marker of self and potent ‘don’t eat me’ inhibitory signal against macrophage phagocytosis, and antibody labeling attenuates its inhibitory effects. Many cancer cells express high levels of CD47 to evade phagocytosis by the innate immune system, including cultured Jurkat T cells, which we chose for initial testing. Macrophages that only trogocytose would be marked by only an AlexaFluor647 signal, while those that phagocytose (as well as that phagocytose and trogocytose) would be marked by both an internalized pHrodo signal and AlexaFluor647 signal.

When we mix target Jurkat T cells with RAW 264.7 macrophage-like cells, we observe two clear phenotypes by fluorescence microscopy. Some macrophages fully engulf the labeled Jurkat T cells with surface antibody signals and volume signals observed to be colocalized in the phagolysosome of the macrophage, which are labeled with CellTracker Green CMFDA (Fig. 1A, bottom panel). Other macrophages internalize small ‘bites’ from the surface of Jurkat T cells that are positive for the anti-CD47 antibody surface signal but negative for the volume marker, pHrodo (Fig. 1A, top panel). From fluorescence microscopy images of macrophages incubated with opsonized Jurkat T cells for one hour, we found that a large proportion of macrophages nibbled target cells, while full phagocytosis events were quite rare (Fig. S2).

**Figure 1.**
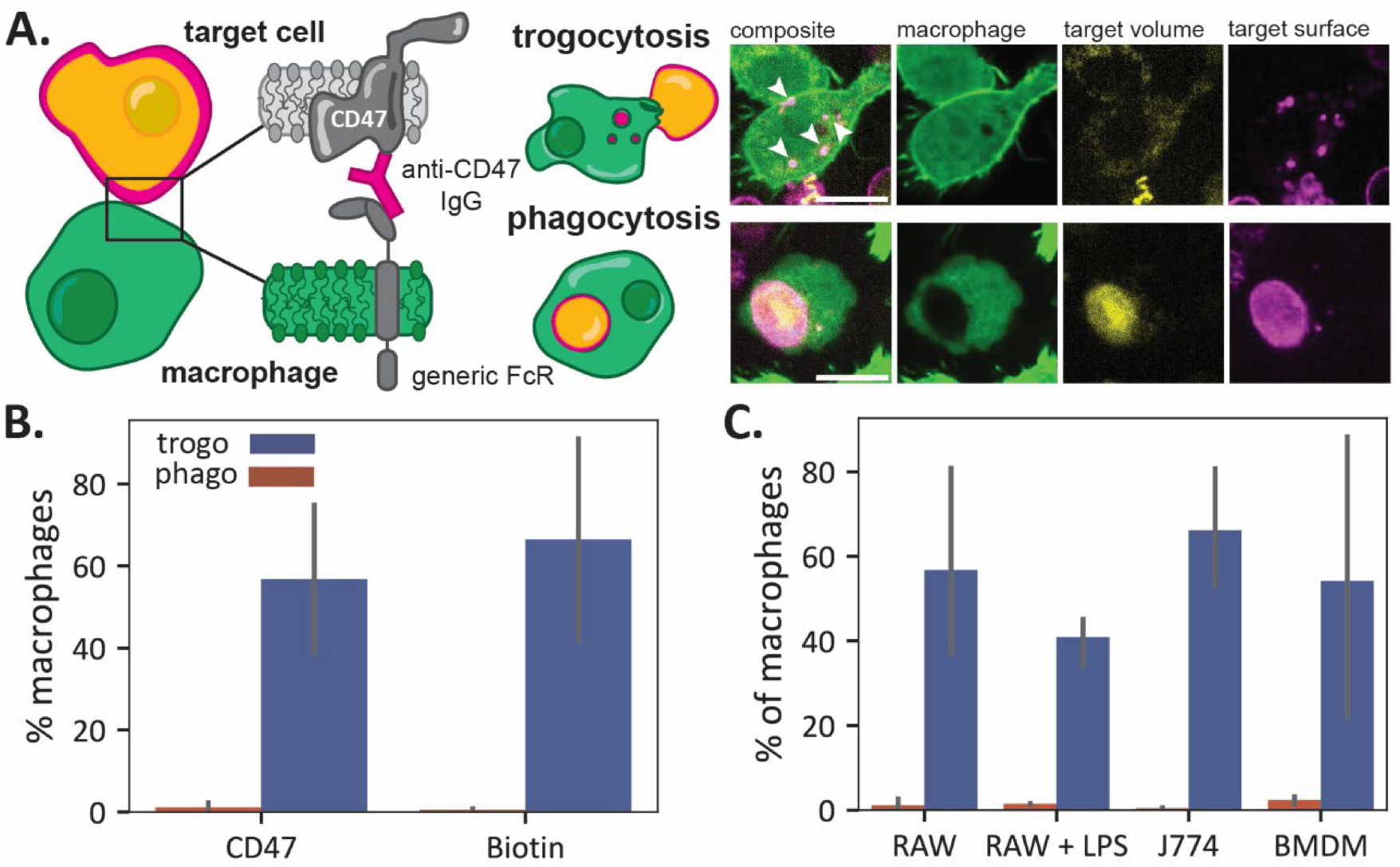
Macrophages trogocytose and phagocytose Jurkat T cells. **A.** Jurkat T cells are labeled with pH-sensitive fluorophore pHrodo and fluorescent (AlexaFluor647) anti-CD47, which binds to the FcR on CellTracker Green CMFDA labeled macrophages, initiating trogocytosis or phagocytosis. Trogocytosis is characterized by internalization of small, punctate ‘bites’ (marked with white arrows) that are positive for the anti-CD47 signal only. Phagocytosis is characterized by colocalization of pHrodo and antibody signal in the phagolysosome of macrophages. The scale bars are 10 μm. **B.** Trogocytic and phagocytic efficiency of RAWs challenged with Jurkat T cells for one hour, opsonized with anti-CD47 and anti-biotin, respectively. **C.** Trogocytic and phagocytic efficiency of RAW 264.7 cells, BMDMs, J774a.1 cells, and LPS-stimulated RAW 264.7 cells challenged with anti-CD47 opsonized Jurkat T cells. Error bars represent the standard deviation between three biological replicates.

To test this observation with larger populations, we developed a flow cytometry assay to quantify trogocytic and phagocytic efficiency. We mixed cytosol-labeled macrophages with surface-and-volume labeled target cells and analyzed the populations of macrophages that contained both target surface and volume signals (indicating phagocytosis), only the surface signal (indicating trogocytosis only), or neither signal. We define phagocytic and trogocytic efficiency as the percentage of the total number of macrophages that have undergone phagocytosis or trogocytosis, respectively.

Consistent with the microscopy experiments, we find that trogocytic events are significantly more frequent than phagocytic events when target cells are opsonized with anti-CD47 (Fig. 1B). We estimate that trogocytosis is ∼3-4x more frequent than phagocytosis in our 1-hour experiments with a 1:1 ratio of macrophages to target cells.

To test whether trogocytosis was dependent on CD47 labeling or could occur with labeling of other cell surface proteins, we non-specifically biotinylated the primary amines of proteins on the surface of target cells using an NHS-ester biotin compound. We then opsonized target cells with anti-biotin, labeled them, and mixed them with macrophages. We found that trogocytic efficiency was nearly identical to that for the anti-CD47 labeling at the same antibody concentration, indicating that trogocytosis is not dependent on the surface molecule labeled (Fig. 1B).

To ensure that the trogocytosis we measure is dependent on FcR binding to the antibody, we measured the trogocytic and phagocytic efficiency of macrophages first treated with an FcR-blocking antibody (mouse anti-CD16/32). Both trogocytosis and phagocytosis are inhibited when the macrophage FcR is blocked by 0.04 μM of antibody (Fig. S3), confirming that our trogocytosis experiments are dependent on FcR engagement.

We next explored whether the trogocytosis we observe is unique to macrophage-like RAW 264.7 cells, which are convenient to work with but have known differences from primary mouse macrophages and other macrophage cell lines^17,18^. We quantified trogocytic and phagocytic efficiency of opsonized Jurkat T cells incubated with primary bone marrow derived macrophages (BMDMs), LPS-stimulated RAW 264.7 cells, and another immortalized mouse macrophage cell line, J774a.1 cells. We observed a similar trogocytic efficiency across all four macrophage cell lines (Fig. 1C). The primary difference among the macrophages is their phagocytic efficiency, with the BMDMs and LPS-stimulated RAWs exhibiting the most phagocytosis, as expected. This shows that both the ability to trogocytose and the trogocytic efficiency is not strongly affected by macrophage cell type.

### Trogocytosis differs for target cells depending on their cortical tension

While different macrophages trogocytose the same target cell with similar efficiency, does the same macrophage trogocytose different target cells with similar efficiency? To test this, we used a panel of cultured cells derived from blood cancers, namely Jurkat T cells, HL60s, and Raji Bs. We opsonized the target cells with anti-CD47, which is highly expressed on all three cell types (Fig. S4) and incubated them with RAW 264.7 macrophages in separate experiments. Interestingly, we see large differences in trogocytic and phagocytic efficiency for the different target cell lines, with the trogocytic efficiency of HL60s and Raji Bs at only ∼20%, compared to ∼70% for Jurkats (Fig. 2A). Notably, phagocytic efficiency is greater for HL60s and Raji Bs than for Jurkats. Since Jurkat T cells have approximately 2.2x more CD47 on their cell surface than HL60s and Raji B cells (Fig. S4), differences in trogocytic efficiency could be due to differences in antibody surface density. We altered antibody solution concentrations to achieve the same surface density of anti-CD47 across cell types (Fig. 2B) and we tested non-specific labeling of the target cells by biotinylating and opsonizing them with anti-biotin (Fig. 2C). In both cases, we observe that trogocytic efficiency is highest in Jurkats, while phagocytic efficiency is highest in HL60s.

**Figure 2.**
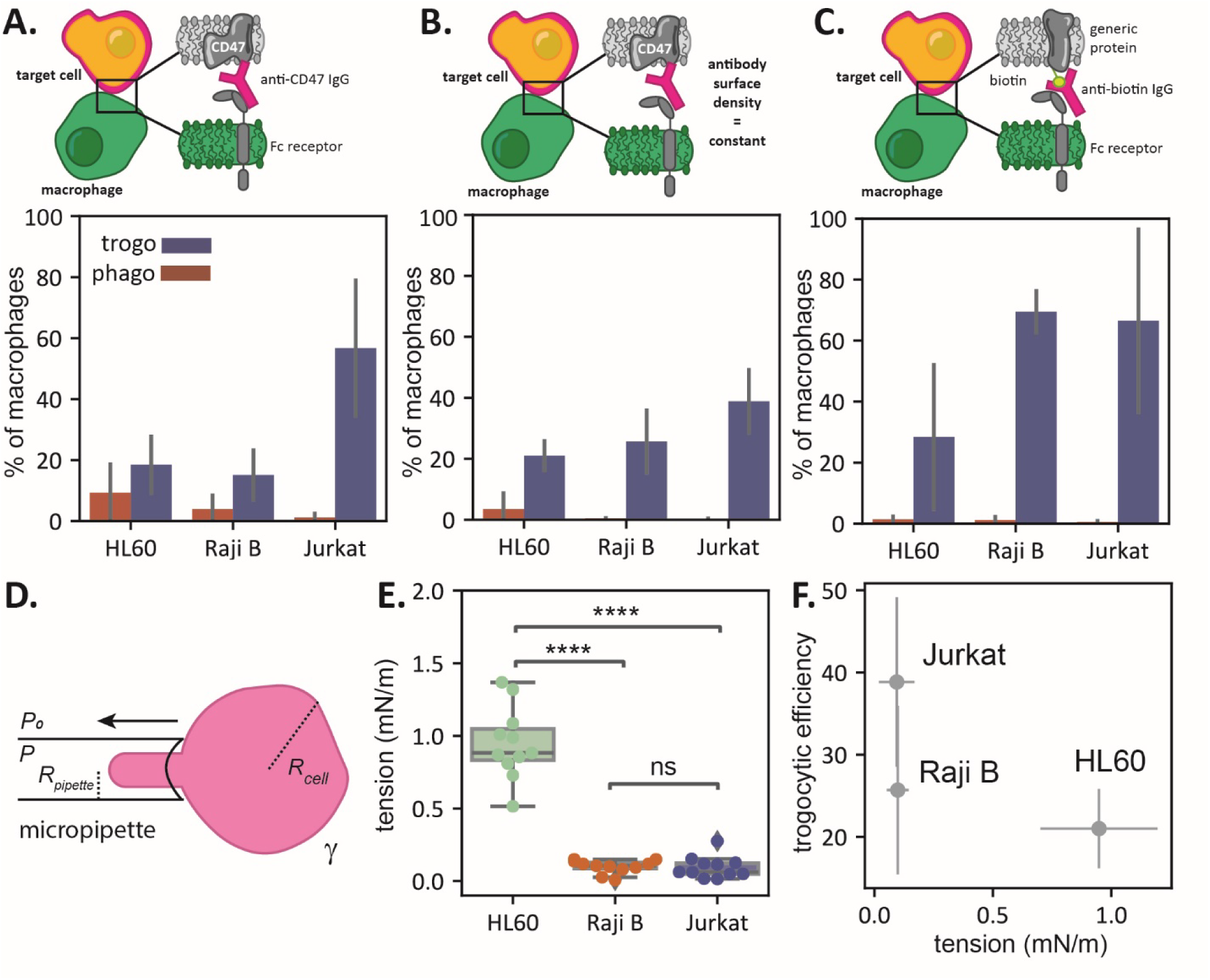
Trogocytosis depends on cell cortical tension. **A.** Phagocytic and trogocytic efficiency of RAWs challenged with Jurkat T cells, Raji B cells, and HL60s opsonized with anti-CD47 at a solution concentration of 0.04 μM. **B.** Phagocytic and trogocytic efficiency of RAWs challenged with Jurkat T cells, Raji B cells, and HL60s opsonized with anti-CD47 solution matched to achieve equivalent surface coverage of antibody (300-500 antibodies/μm^2^). **C.** Phagocytic and trogocytic efficiency of RAWs challenged with Jurkat T cells, Raji B cells, and HL60s with surface proteins non-specifically labeled by NHS-biotin and then opsonized with anti-biotin at a solution concentration of 0.04 μM. **D.** Micropipette aspiration set up to measure cell tension. We measure *Rcell*, *Rpipette*, and *P* to calculate the surface tension, γ. **E.** Tension of Raji B cells, HL60s, and Jurkat T cells. There is a significant difference (computed via one-way ANOVA followed by a Tukey’s pairwise test) between the tension of HL60s and Raji Bs and Jurkat T cells. **F.** The trogocytic efficiency of target cells from Fig. 2b is negatively correlated with target cell tension (Pearson’s correlation coefficient of -0.71).

Since trogocytosis involves pinching off small (<1 μm) bits of membrane, one potentially relevant difference between the target cells could be the deformability of their cell surface. This deformability is dependent not only on membrane tension but also on the strength and abundance of linkages between the membrane and underlying cytoskeleton. Together, the resistance to deformation can be captured by ‘cortical tension’, often measured with micropipette aspiration^19–21^. We quantified the cortical tension of antibody-labeled target cells using a home-built micropipette aspiration system with an in-line pressure sensor. Using glass pipettes with an inner diameter of ∼5 μm, we applied suction to individual cells until a ‘tongue’ is pulled into the syringe to a consistent length of ∼1.5 μm (Fig. 2D). We then quantified cortical tension of the cell using the Young- Laplace law,

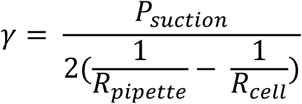

where *ΔPsuction, Rpipette*, and *Rcell* are the suction pressure, the pipette radius, and the cell radius.

We find that the cortical tension of HL60s, which are phagocytosed more than the other cells, is significantly higher than that of the Jurkat T cells and Raji B cells, which are trogocytosed more than HL60s (Fig. 2E). If we take the average trogocytic efficiency of target cells with matched anti-CD47 surface densities, the trogocytic efficiency for target cells decreases significantly (Pearson’s correlation coefficient of -0.71) with increasing cell tension (Fig. 2F).

### Membrane tension of a cell mimic drives trogocytosis over phagocytosis

The cortical tension we measured for cells is a combination of membrane tension and interactions with the cortical cytoskeleton. We wondered whether membrane tension alone could be used to alter macrophage trogocytosis, independent of any cytoskeletal interactions. To test this, we created target particles with only membrane using giant unilamellar vesicles (GUVs) composed of palmitoyl oleoyl phosphatidylcholine (POPC), 1 mol% biotin dioleoyl phosphatidylethanolamine (biotin-DOPE), and 0.5 mol% lissamine rhodamine phosphatidylethanolamine (liss rho PE). The GUVs were formed in a sucrose medium that was osmotically tuned using an osmometer to cause the GUVs to sink but remain isotonic in the cell culture media. To obtain GUVs of a consistent size and avoid the confounding effects of small vesicles formed at the same time as the GUVs, we developed a home-built differential density column (SeparatorMax 3000, Fig. S1) to separate the GUVs based on size. We collected fractions of GUVs in the size range of 5-20 μm in diameter and opsonized them with fluorescent anti-biotin at a surface density of ∼400 antibodies/μm^2^ for a 10 μm GUV.

We incubated opsonized GUVs with macrophages and observed by confocal microscopy that macrophages can both fully engulf GUVs and partially nibble them (Fig. 3A). When a GUV is trogocytosed, we observe colocalization of the lipid dye and the fluorescent antibody in trogocytic bites, indicating that the macrophages extract small patches of membrane from GUVs rather than detach the anti-biotin AF647 antibody. These initial experiments showed that antibody-opsonized GUVs can be both trogocytosed and phagocytosed, similar to target cells.

**Figure 3.**
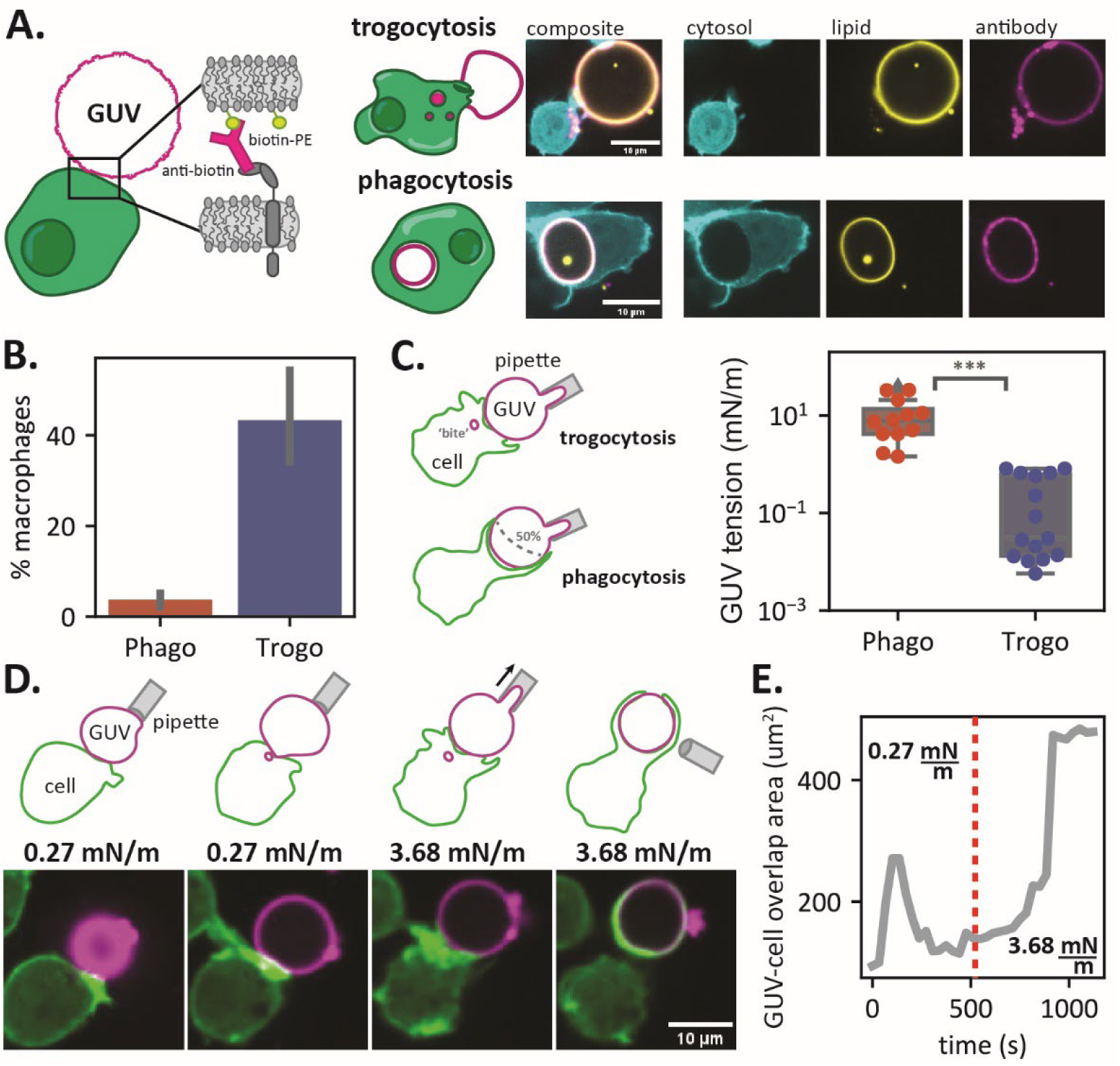
Macrophages trogocytose GUVs. **A.** GUVs composed of POPC, biotin-DOPE, and liss rho PE are opsonized with AlexaFluor647 anti-biotin and mixed with macrophages. When GUVs are phagocytosed, the perimeter of a circular GUV can be observed within the macrophage phagolysosome (bottom) and when GUVs are trogocytosed, punctate ‘bites’ cab be observed within the macrophage (top). **B.** GUVs are trogocytosed more frequently than phagocytosed by macrophages. **C.** GUVs that are phagocytosed (determined by >50% surface coverage of the GUV by macrophages phagocytic extensions) are in a high-tension regime (>1 mN/m), and GUVs that are trogocytosed (determined by the presence of trogocytic ‘bites’ in the macrophage) are in a low-tension regime (< 1 mN/m). Significance was determined via a student’s T test. **D.** Schematic and micrographs of a GUV under low tension getting trogocytosed by a macrophage, followed by application of suction and an increase in tension, leading to full engulfment of the GUV by the macrophage. **E.** Time trace of the overlap area between the macrophage channel (green) and the GUV channel (magenta). The dashed red line indicates when suction was applied to the GUV and tension was increased from 0.27 mN/m (trogocytosis regime) to 3.68 mN/m (phagocytosis regime).

To quantify the trogocytic and phagocytic efficiency of macrophages incubated with GUVs, we loaded dark GUVs with a soluble rhodamine fluorophore and opsonized them with AlexaFluor647 anti-biotin. After 30 minutes of co-incubation with macrophages (from microscopy experiments we observed that macrophage phagocytosis and trogocytosis of GUVs is much more rapid than of target cells), we analyzed the macrophages via flow cytometry and found that trogocytosis is a much more common event than phagocytosis, consistent with the target cells tested (Fig. 3B). We might have expected that all GUVs would either by trogocytosed or phagocytosed since they all have the same lipid composition and vary only minimally in size. However, something must be different – even in this minimal system – that is sufficient to drive trogocytosis rather than phagocytosis.

Interestingly, GUVs created via electroformation are formed with a wide range of membrane tensions, even when they are ‘isotonic’ (Fig. S5). To directly investigate the role of membrane tension in trogocytosis, we first used micropipette aspiration to measure the membrane tension of a population of GUVs. We incubated GUVs with macrophages in an imaging dish, allowed GUVs to settle and iteract with the macrophages for 10 minutes, and then individually measured the membrane tension of GUVs that were being either phagocytosed or trogocytosed. We identified phagocytic events as GUVs whose surface was wrapped >50% by macrophage phagocytic extensions and trogocytic events as GUVs connected to macrophages in which small ‘bites’ were internalized in the macrophage. We found two distinct tension regimes: At tensions < 1 ± 0.33 mN/m, GUVs are exclusively trogocytosed, while at tensions > 1 ± 11.2 mN/m GUVs are phagocytosed (Fig. 3C).

These measurements show that membrane tension is a sufficient signal to drive macrophage trogocytosis or phagocytosis of GUVs. However, is that decision made at initial contact and activation, or does the macrophage dynamically sense membrane tension and alter its decision? To address this, we tested the response of macrophages to dynamic changes in membrane tension with the micropipette aspiration system and simultaneous confocal microscopy. We added opsonized GUVs to one side of an imaging chamber and selected a GUV with a low membrane tension (0.27 mN/m; in the trogocytosis regime). Upon presenting the GUV to a macrophage, we observed the macrophage form filopodial extensions, but then quickly retract and begin internalizing small ‘bites’ of a GUV membrane. Using the micropipette, we then applied suction to the GUV, increasing membrane tension above the threshold for phagocytosis (3.68 mN/m; in the phagocytosis regime). Within three minutes, the macrophage rapidly extended a phagocytic cup around the GUV and engulfed it completely within a matter of minutes (Fig. 3D-E; movie S1; movies S2-S3 for more examples). These experiments show that membrane tension alone is a significant driver of trogocytosis in macrophages, and changes in membrane tension can act as a switch between phagocytosis and trogocytosis.

### Increasing tension in cultured cells suppresses trogocytosis

Our experiments with cultured cells and GUVs indicate that deformability of the target surface is sufficient to control whether the target is phagocytosed or trogocytosed. This raises the possibility that intentionally stiffening the cortex of cells could suppress trogocytosis and promote phagocytosis. Previous work has shown that gentle glutaraldehyde fixation leads to measurable differences in corticall tension^22^, with concentrations of glutaraldehyde below 0.01% leaving cells viable. We treated opsonized target cells with either 0%, 0.002%, or 0.0025% glutaraldehyde for 30 minutes. At these fixation concentrations, anti-CD47 density on treated target cells is approximately the same as untreated cells (300-500 ± 100 antibodies/μm^2^).

To confirm the effects of gentle fixation on cortical tension, we measured treated target cells using micropipette aspiration. We chose HL60s and Jurkats because they engage in the least and most trogocytosis, respectively. Cortical tensions for treated cells increased above the tension regime we measured for trogocytosis of GUVs, especially for Jurkats and HL60s treated with 0.0025% glutaraldehyde (Fig. 4A). To quantify the trogocytic efficiency of macrophages with the opsonized and stiffened target cells, we washed and incubated them with macrophages for one hour and then analyzed them with flow cytometry. We found that trogocytosis was almost completely eliminated for cells treated with 0.002% glutaraldehyde or higher (Fig. 4B). In contrast, phagocytosis increased for macrophages challenged with cells treated with glutaraldehyde (Fig. S6).

**Figure 4.**
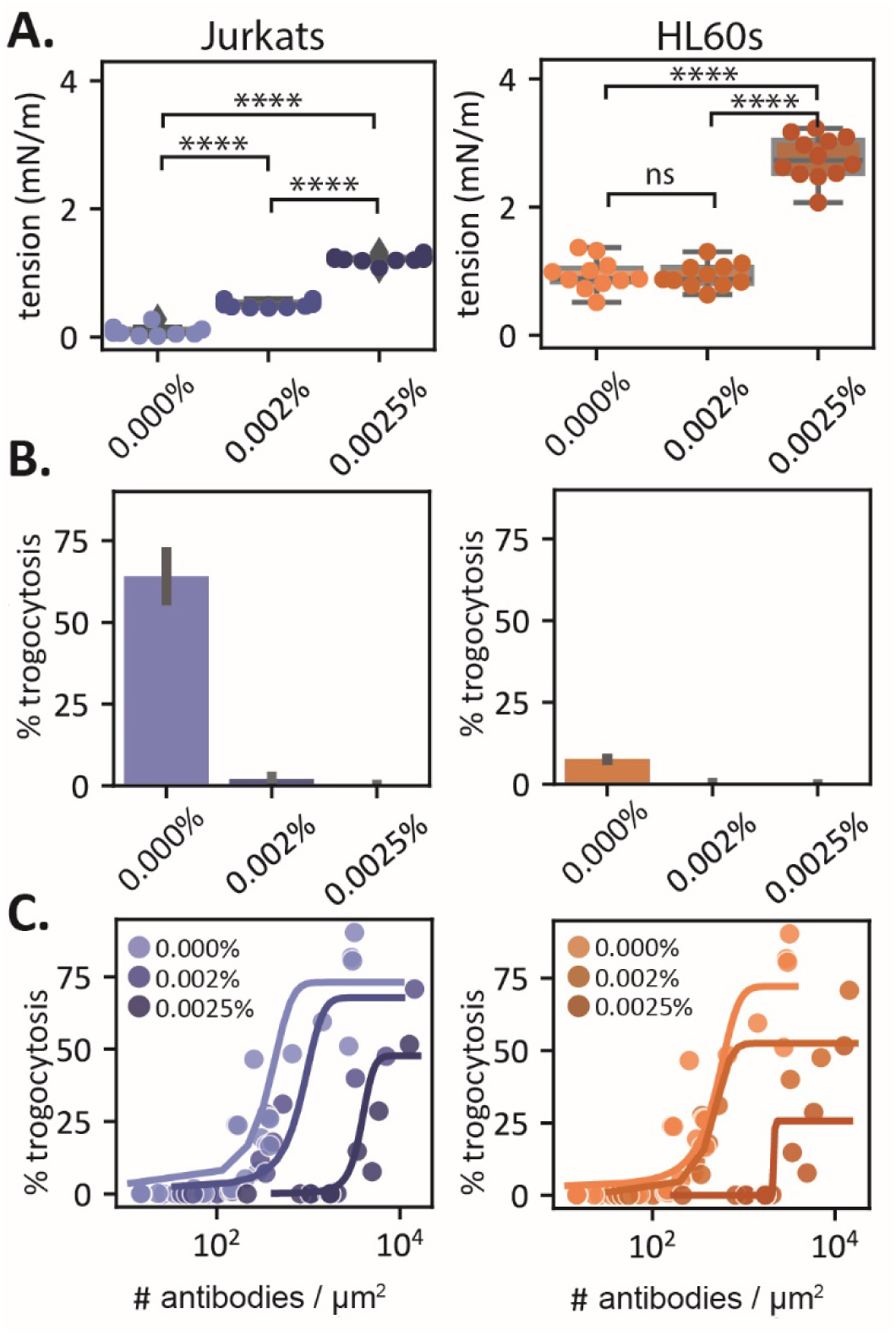
Increasing cell tension via gentle fixation suppresses trogocytosis. **A.** Cell tension measurements for Jurkats and HL60s treated with 0%, 0.02%, and 0.0025% glutaraldehyde. Significance was determined via a one-way ANOVA followed by a Tukey’s pairwise test. **B.** Trogocytic efficiency of macrophages challenged with glutaraldehyde-treated target cells. **C.** Trogocytic efficiency of macrophages challenged glutaraldehyde-treated target cells with a surface density titration of anti-CD47. Sigmoid curves (described in the text) fit to points to guide the eye.

Since we previously observed that Jurkats, before antibody surface density matching with HL60s and Raji Bs, had greater trogocytic efficiency with greater antibody coverage, we wondered if we could rescue trogocytosis of the stiffened cells by increasing antibody surface density. When we titrated antibody surface density for gently fixed Jurkats and HL60s, we found that trogocytic efficiency was indeed increased at the highest antibody densities (Fig. 4C). Interestingly, trogocytic efficiency is rescued at much higher concentrations of antibody than for wild type target cells. Not only does it take more antibody to induce trogocytosis of gently fixed cells, the trogocytic efficiency is never greater than what was achieved for wild type target cells at lower antibody densities (Fig. S7). This suggests that we are unable to coat the surface sufficiently to rescue the full trogocytic efficiency of a soft wildtype target.

### A mechanical model captures the relationship between cell tensions and antibody surface density in trogocytosis

Our experiments indicate both cortical tension and antibody density play important roles in driving trogocytosis of cellular targets. How might the two interact to control interface mechanics between the macrophage and target cell? To gain intuition, we develop a simple scaling law for the interfacial mechanics. We assume the target antibody binds to macrophage FcRs at a high enough concentration to trigger downstream signaling machinery that leads to cytoskeleton-mediated active stresses at the interface (Fig. 5A). Based on prior experimental measurements these stresses take the form of both extensile and compressive normal stresses on the target membrane^3^. We further assume that these stresses scale with the antibody density on the target (Supplementary Text). These normal stresses will tend to deform the interface, and this deformation is resisted by the target cortical tension. Balancing the forces due to active stresses and cortical tension at the interface, one gets:

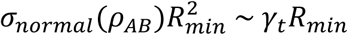

This gives the length scale:

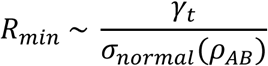

**Figure 5.**
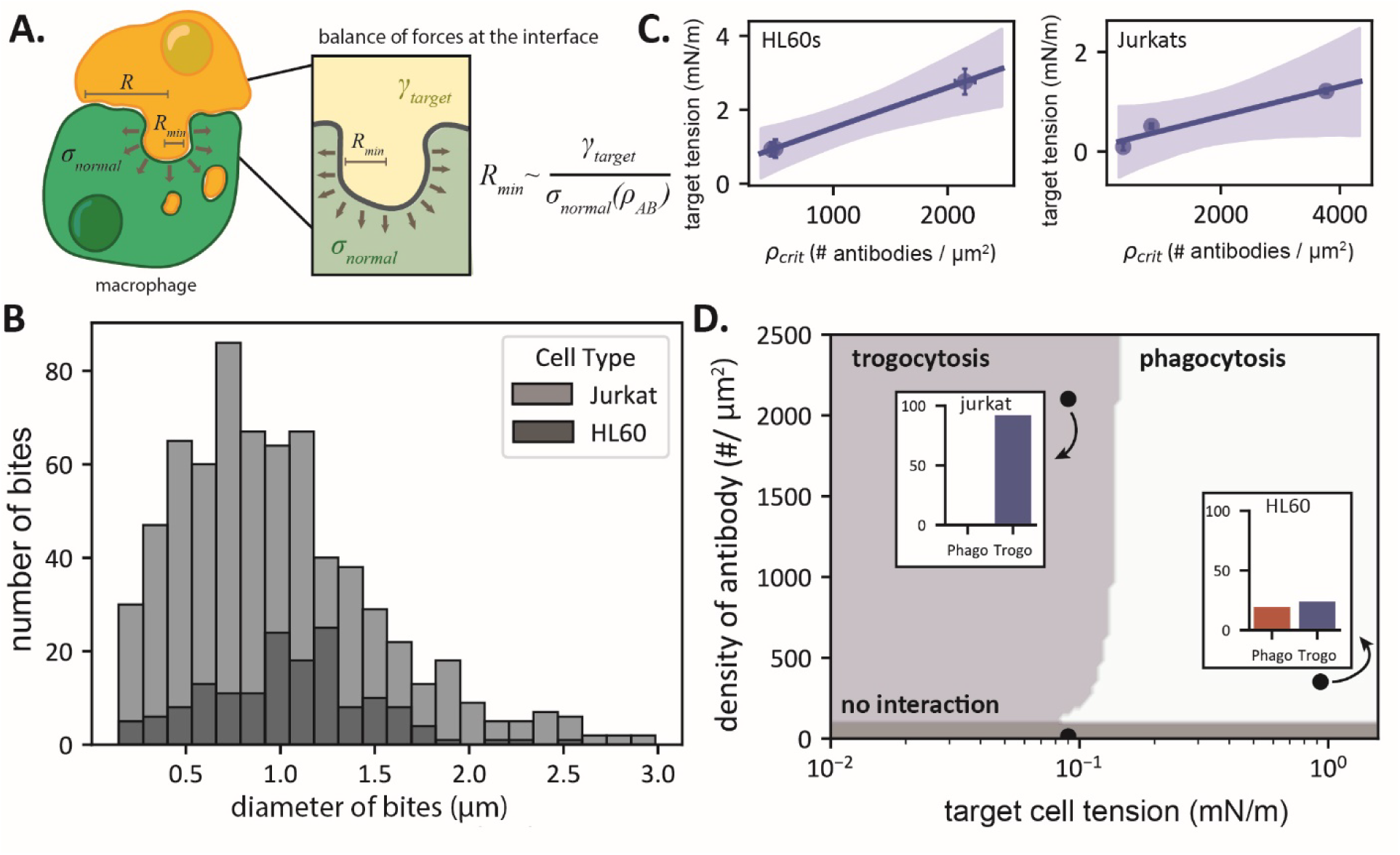
Mechanical scaling law describing macrophage trogocytosis. **A.** A cartoon model of the macrophage-target interface. **B.** Distribution of trogocytic ‘bite’ sizes for macrophages co-cultured with Jurkats (light grey) or HL60s (dark grey) measured from confocal images of macrophages post co-culture. **C.** ρcrit scales with the cell tension for treated HL60s and Jurkats. Shaded area corresponds to the 95% confidence interval of the linear fit, computed via bootstrap resampling. Uncertainty on the points in the x-axis represents 95% confidence intervals for the computed inflection point of the sigmoid fits and the standard deviation across cell tension measurements on the y-axis. **D.** Approximate phase diagram of macrophage behaviors at different antibody densities and effective membrane tensions based on the scaling law for bite-size. Points correspond to individual measurements at a particular antibody density and cell tension and their corresponding phagocytic and trogocytic efficiencies. Targets in this case are either Jurkat T cells or HL60s.

where γt is the target cortical tension and σnormal(ρAB) is the antibody-dependent normal stress at the interface. This scaling corresponds to the minimum length scale of local membrane deformation or bites, since below this scale tension dominates the effects of active stresses and damps out active fluctuations. A similar length scale can be derived by considering the balance between active stresses and membrane bending (Supplementary Text), which is typically smaller than *Rmin* for physiologically relevant values of tension and membrane bending stiffness.

Importantly, this scaling implies that increasing antibody density or decreasing target membrane tension has an equivalent effect on setting the minimum length scale of target membrane deformations. For small antibody density or large tensions, the active stresses are weak compared to those due to tension, and the minimum scale of deformation due to active stress becomes comparable or larger than the interface size *R* (*Rmin* >> *R*). In this case, the macrophage is incapable of pinching off bites smaller than the interface and would completely engulf the target (i.e., phagocytosis). In contrast, for small tensions or large antibody densities, the active stresses can locally overwhelm membrane tension and pinch-off bites smaller than the interface size (*Rmin* << *R*), leading to trogocytosis. Below a second, lower critical antibody density, one expects no engagement and neither behavior is observed. Thus, one can also interpret this scaling as a critical antibody density or tension that separates the distinct behaviors of phagocytosis and trogocytosis.

If we take our experimentally measured values for γt ∼ 10^-2^ – 10^-1^ mN/m and the previously measured traction forces that give normal stresses of order 100 Pa ^3^, our scaling predicts a deformation size *Rmin* of 0.1 – 1 μm. Interestingly, this is consistent with the scale of trogocytic ‘bites’ measured through confocal microscopy, which are on the scale of 1 μm for macrophages incubated with Jurkat T cells and HL60s (Fig. 5B). This suggests that the macrophage can pinch off bites of a size close to the minimum scale of interfacial deformation.

Our scaling also predicts that the critical antibody density that separates phagocytic and trogocytic behaviors in experiments must be proportional to the target cell cortical tension. To test this, we fit the sigmoidal curve below to the data in Fig. 4c and extracted the inflection point of the curve as the critical antibody density, *ρcrit*:

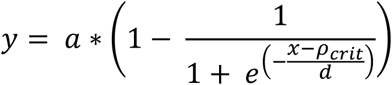

For both HL60s and Jurkats, we find that the density of antibody necessary to initiate trogocytosis indeed increases with cortical tension (Fig. 5C).

From the scaling relationship described above and assuming an interface size R of ∼1 μm, we predict a tension dependent switch in macrophage behavior at a tension around 0.1 – 1 mN/M for the range of normal stresses measured in experiments (50-150 Pa)^3^. Based on this prediction, we can construct an approximate phase diagram that predicts the macrophage behavior as a function of antibody density and cortical tension (Fig. 5D). Our experimental data for Jurkat and HL60 cell lines are quantitatively consistent with this phage diagram, with cortical tension predicted to have a more significant effect than antibody density.

## Discussion

Immune cells, including macrophages, are in regular physical contact with cells around them, both ‘self’ and foreign. It is increasingly clear that the physical properties of target cells^3,23–28^ and their surface molecules^29–33^ can influence the response of immune cells during phagocytosis, T cell activation, dendritic cell antigen presentation, and B cell activation. In this work, we show that the cortical tension of target cells is a key regulator of macrophage decision making between phagocytosis and trogocytosis.

Using fluorescence microscopy and flow cytometry, we demonstrate that trogocytosis is consistently more common than phagocytosis. Trogocytic efficiency with Jurkat T cells is regularly between 60-80% of events, while the highest phagocytic efficiency we measured for HL60s is only ∼20%. Perhaps this is not surprising given that a target cell can be trogocytosed by many cells but phagocytosed by only one. Still, if trogocytosis is so ubiquitous, why is it rarely quantified in studies of macrophage effector functions? It is possible that trogocytosis is happening in most phagocytosis assays involving soft targets (i.e., cells) but is missed if target particles/cells are not labeled with a membrane marker. In this study, we can only ‘see’ trogocytosis if the membrane of targets is sufficiently labeled. Indeed, in studies where the membrane is labeled, trogocytosis is observed and often characterized as a negative outcome in phagocytosis assays^5,34^.

To address the fundamental question of what governs macrophage trogocytosis, we explored the conditions that lead to either phagocytosis or trogocytosis. We observed that trogocytosis and phagocytosis both require FcR engagement with antibodies on the surface of the target cell, and that both require a certain threshold antibody density (∼10^2^ antibodies/μm^2^) to occur. Importantly, the identity of the opsonized surface protein does not appear to matter, while the total surface density of antibody influences the degree of trogocytosis for a given cell type.

In our previous experiments using lipid-coated glass beads as targets, macrophages readily phagocytosed but never seemed to trogocytose the membranes on the target beads. This led us to test whether macrophages could phagocytose free-floating and deformable GUVs composed of the same lipids typically found in lipid-coated glass bead assays. We saw that when a target particle is capable of being deformed, as is the case for GUVs, trogocytosis is readily observed and much more frequent than phagocytosis. Our surface tension measurements of GUVs show that membrane tension is a key regulator of trogocytosis by macrophages, with two distinct membrane tension regimes for phagocytosis and trogocytosis. By dynamically switching a GUV from one to the other, we observe that macrophages rapidly transition from trogocytosis to phagocytosis.

The pervasiveness of trogocytosis in antibody-mediated interactions with soft targets like cells and GUVs is not entirely surprising if we consider the cellular materials required to build a phagocytic cup. To engulf a large particle, like a cancer cell, a macrophage must assemble enough actin cytoskeleton and incorporate sufficient excess membrane to fully wrap and exert force upon an object that is roughly the same size as the macrophage. In contrast, trogocytosis requires significantly less cellular machinery than phagocytosis. For example, we measured the distribution of trogocytic ‘bite’ sizes for HL60s and Jurkats and on average, each ‘bite’ is ∼ 1 μm in diameter, which is considerably easier to engulf than cell-sized objects.

With enough FcRs clustered to locally initiate branched actin network assembly, a soft and deformable target membrane could easily be pinched off at the macrophage-target interface. Indeed, in high-resolution fluorescence microscopy videos of macrophages with HL60s, we see multiple ‘bites’ being extracted from the HL60 by the macrophage along the surface of the cell-cell interface (Movie S4). This observation hints at the intriguing possibility that macrophages are constantly surveying and extracting small bites from their surroundings. Only if they encounter a foreign pathogen, such as a stiff bacterial cell (100-200 MPa for *Pseudomonas aeruginosa*^35^), do they opt for phagocytosis.

Our identification of cortical stiffness as a key factor in the decision between trogocytosis and phagocytosis raises the possibility that it could be beneficial for tumor cells to increase their deformability to reduce the possibility of phagocytic interactions with macrophages. Some tumor cells are known to have reduced stiffness compared to their health counterparts^36,37^, and immunotherapeutic antibodies targeting tumor cells have been observed to be trogocytosed^10^, removing the antibody and leaving the tumor cells alive and less visible to the immune system. Therapies that stiffen the cortex of target cells could have the benefit of promoting phagocytosis and limiting trogocytosis.

## Resource Availability

Lead contact:

Requests for further information and resources should be directed to and will be fulfilled by the lead contact, Daniel A. Fletcher.

Materials, data, and code availability:

This study did not generate new unique reagents. Original code to generate figures in the paper is available at: https://github.com/fletchlab-git/Trogocytosis. Datasets used to generate figures in the paper are available at: DOI 10.17605/OSF.IO/6FP4V

## Author Contributions

C.E.C., A.C., D.K., and D.A.F. designed the research. C.E.C., A.C., N.R.M., and L.B., performed micropipette aspiration experiments. C.E.C., N.R.M., and L.B., performed trogocytosis/phagocytosis flow cytometry experiments with cultured target cells. C.E.C. performed trogocytosis/phagocytosis flow cytometry and microscopy experiments with GUVs. D.K. designed and fabricated the SeparatorMax 3000. D.K. derived the scaling law. C.E.C. analyzed data. C.E.C., A.C., D.K., and D.A.F. wrote the paper.

## Acknowledgements

We thank all of the members of the Fletcher lab for discussions, assistance, and advice. We thank the Woods Hole Marine Biological Laboratory Physiology Summer Course for providing the brilliant students Donovan Phua and Austin J. Graham, both of whom contributed to this research at different stages. We also thank FIP (France Inter Paris) Radio for inspiration during long micropipette sessions at the microscope. This work was supported by an NSF Center for Cellular Construction DBI-1548297 (D.A.F.) and an NIH R01 GM134137 (D.A.F). C.E.C. was supported by a James S McDonnell Foundation Postdoctoral Fellowship. A.C. was supported by an EMBO Postdoctoral Fellowship. D.K. was supported by a Schmidt Science Fellowship in partnership with the Rhodes Trust. D.K. is also supported by a Burroughs Wellcome Career Award at the Scientific Interface. N.R.M. was supported by a Cancer Research Institute Postdoctoral Fellowship. L.B. was supported by a Stichting Fonds Doctor Catharine van Tussenbroek Travel Grant. D.A.F. is a Chan Zuckerberg Biohub Investigator.

## Supplemental Information

Document S1. Figures S1-S8 and Supplemental Text

Movie S1. Applying suction to a GUV switches a macrophage from trogocytosis to phagocytosis. Related to Figure 3.

Movie S2. Applying suction to a GUV switches a macrophage from trogocytosis to phagocytosis. Related to Figure 3.

Movie S3. Releasing suction on a GUV switches a macrophage from phagocytosis to trogocytosis. Related to Figure 3.

Movie S4. Volumetric movie of a macrophage trogocytosing an HL60. Related to Figure S5.

