## Supplemental Information for "Target cell tension regulates macrophage trogocytosis"

**CONTENTS**

**Figure S1.** Design of the SeparatorMax3000.

**Figure S2.** Fluorescence micrographs of HL60s co-cultured with RAW 264.7 cells.

**Figure S3.** Titration of anti-CD16/32 (FcBlock) on macrophages.

**Figure S4.** CD47 expression on target cells.

**Figure S5.** Membrane tension of GUVs in an isotonic solution.

**Figure S6.** Macrophage phagocytosis of glutaraldehyde-treated target cells.

**Figure S7.** Trogocytosis of Jurkat T cells with increasing antibody surface density.

**Figure S8.** Schematic of force balance in the proposed scaling law.

**Movie S1.** Applying suction to a GUV switches a macrophage from trogocytosis to phagocytosis.

**Movie S2.** Applying suction to a GUV switches a macrophage from trogocytosis to phagocytosis.

**Movie S3.** Releasing suction on a GUV switches a macrophage from phagocytosis to trogocytosis.

**Movie S4.** Volumetric movie of a macrophage trogocytosing an HL60.

**Supplementary Text.** Detailed discussion of scaling law derivation.

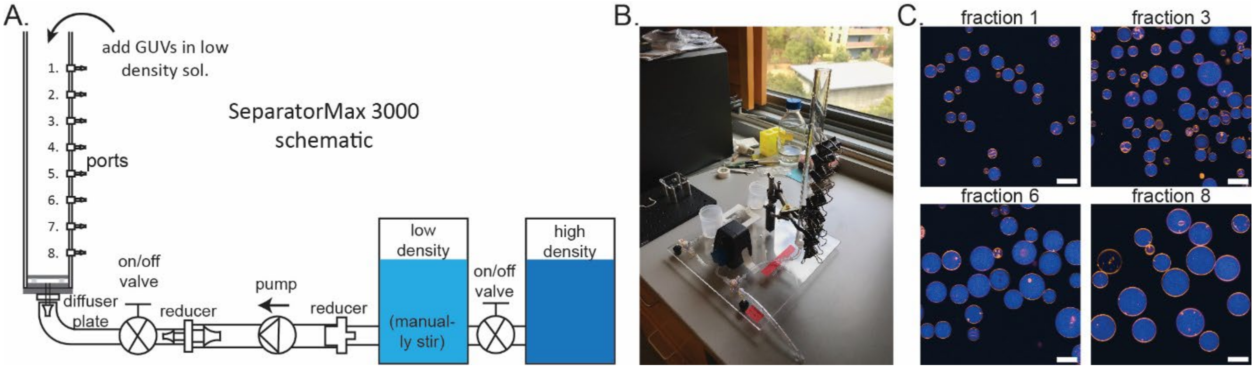


**Figure S1:** Design of the SeparatorMax 3000, a tool for separating GUVs based on size. **A.** Schematic of the SeparatorMax 3000. A primary feature of the design is two different reservoirs of different density solutions. Solutions are pumped into the column while mixing to create a uniform density gradient that opposes convection. GUVs are loaded from the top of the column and allowed to settle for 16-24hrs. GUVs are harvested from different sections of the columns through the outlet ports. **B.** Photo of the SeparatorMax 3000. The ports for retrieving specific fractions are sealed with binder clips until needed. **C.** Confocal fluorescence images of GUVs harvested from different fractions (fraction diameters: one = 14.6 ± 1.6 μm; three = 14.2 ± 5.3 μm; six = 25.9 ± 6.9 μm; eight = 31.1 ± 5.3 μm). GUVs are composed of 99.5 mol% POPC and 0.5 mol% liss rho PE and they are filled with soluble FITC. Scale bars are 25 μm.

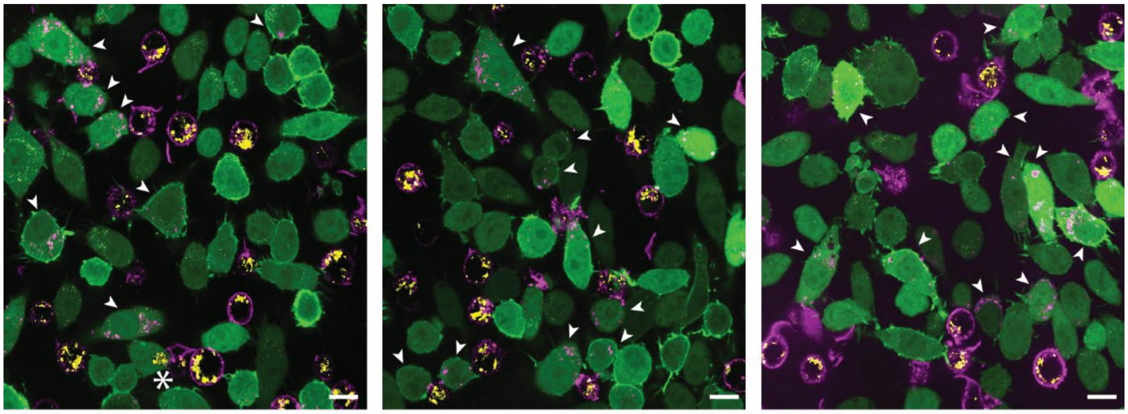

**Figure S2.** Trogocytosis is observed much more frequently than phagocytosis in macrophages. A series of confocal fluorescence images from the same experiment of RAW 264.7 macrophages (green; CellTracker Green CMFDA and LifeAct GFP) incubated with anti-CD47 opsonized Jurkat T cells (magenta and yellow; AlexaFluor 647 anti-CD47 (membrane) and pHrodo (endosomes)). Stars indicate phagocytosis events in which both pHrodo and AF647 signals from Jurkats are present inside a phagosome of a RAW cell. Arrows indicate macrophages that have internalized small trogocytic bites positive only for AF647 from Jurkat T cells.

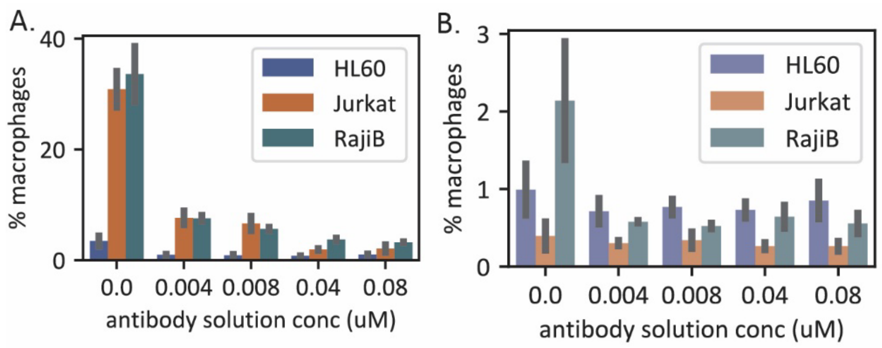


**Figure S3.** Titration of anti-CD16/32 (FcBlock) on macrophages before incubation with anti-CD47 opsonized target cells reduces both trogocytosis and phagocytosis. **A.** Trogocytosis in macrophages is reduced with increasing FcR blocking antibody. **B.** Phagocytosis in macrophages is reduced with increasing FcR blocking antibody.

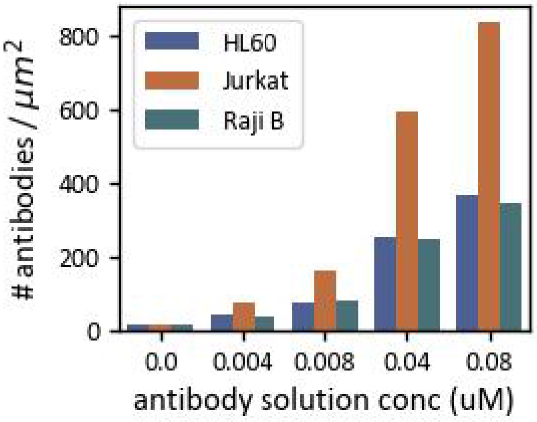


**Figure S4.** CD47 expression on the surface of Jurkat T cells is ~2.2x higher than on HL60s and Raji B cells at a concentration of 0.04 μM.

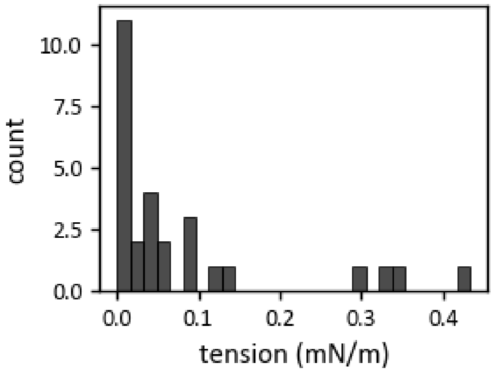


**Figure S5.** Membrane tensions of GUVs in an 'isotonic' solution still vary over an order of magnitude.

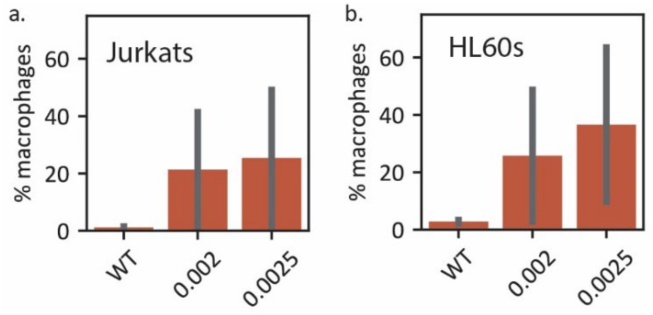


**Figure S6.** Macrophages phagocytose glutaraldehyde-treated target cells. **A.** The amount of macrophages that phagocytose treated Jurkat cells increases with increasing solution concentration of glutaraldehyde. **B.** The amount of macrophages that phagocytose treated HL60s increases with increasing solution concentration of glutaraldehyde.

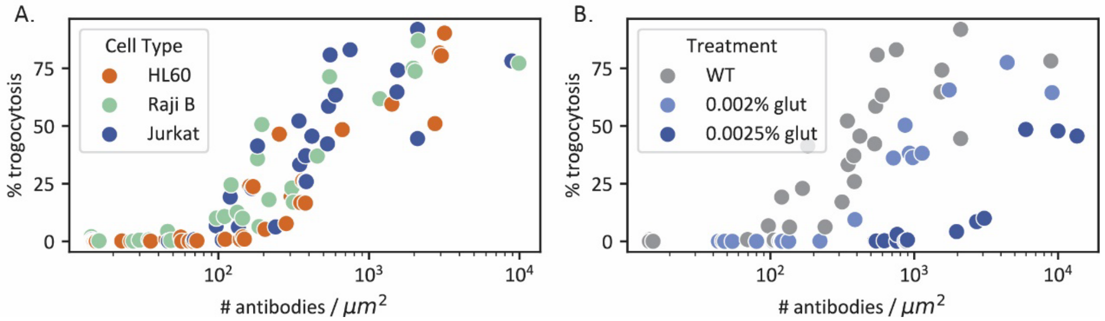


**Figure S7.** Trogocytosis is partially recovered for glutaraldehyde-treated Jurkat cells if the antibody surface density is increased. **A.** The amount of trogocytosis in macrophages increases with surface antibody density for all target cells. **B.** While trogocytosis increases with surface antibody density on glutaraldehyde-treated Jurkats, the amount of trogocytosis is always lower than wildtype cells.

**Movie S1.** Increasing the tension on a GUV switches a macrophage from trogocytosis to phagocytosis. Suction is applied via micropipette aspiration and indicated by the appearance of a white arrow in the movie. The macrophage cytosol is labeled by CellTracker Green CMFDA and the actin is labeled by LifeAct GFP. The GUV is opsonized with fluorescent anti-biotin AlexaFluor 647.

**Movie S2.** Increasing the tension on a GUV switches a macrophage from trogocytosis to phagocytosis. Suction is applied via micropipette aspiration and indicated by the appearance of a white arrow in the movie. The macrophage cytosol is labeled by CellTracker Green CMFDA and the actin is labeled by LifeAct GFP. The GUV is opsonized with fluorescent anti-biotin AlexaFluor 647.

**Movie S3.** Releasing the tension on a GUV switches a macrophage from phagocytosis to trogocytosis. Suction is applied via micropipette aspiration at the beginning of the movie and released part way through. The macrophage cytosol is labeled by CellTracker Green CMFDA and the actin is labeled by LifeAct GFP. The GUV is opsonized with fluorescent anti-biotin AlexaFluor 647.

**Movie S4.** Macrophages consume small fragments from the membrane of HL60s via trogocytosis. The macrophage cytosol is labeled by CellTracker Green CMFDA and the actin is labeled by LifeAct GFP. The Jurkat T cell is opsonized with fluorescent anti-CD47 AlexaFluor 647 and the cytosol is labeled with the pH sensitive dye, pHrodo.

### Supplementary Text

#### Membrane mechanics scaling for trogocytosis

Here we present details and lay out the assumptions that go into deriving the scaling relationship given in the main text. We assume that deformability is mainly controlled by the target properties, though this assumption can be relaxed to include the composite properties of the macrophage and target membranes.

We start by assuming that there is local binding of the target membrane to the macrophage that is FcR-antibody mediated. For a sufficiently high density of antibody, macrophage binding will trigger downstream signaling machinery that leads to cytoskeleton-mediated active stresses at the interface. These are due, in part, to nucleation of branched actin at the region of binding that drives a height-dependent exclusion of tall bystander proteins and their associated inhibitory phosphatases[1]. Prior measurements have shown that these active stresses take the form of compressive and extensile normal stresses that deform the interface[2]. Here we also assume that the magnitude of these stresses is dependent on the antibody density on the target, because more antibody molecules can bind more FcR per unit area and hence drive increased local active stress.

Active deformation of the macrophage is resisted by the cortical tension of the target cell. One can therefore derive a scaling relationship by balancing active stresses and cortical tension.

Consider a local deformation of scale $R_{min}$, due to active stresses (Figure S8). Integrating the effects of active stresses and cortical tension gives:

$$\int_A \sigma_{normal}\, dA \sim \int_S \gamma_t\, ds$$

Where $A \sim R_{min}^2$ is the area over which the normal stresses act, while $S \sim R_{min}$ is the contour length over which tension resists the deformation. A critical scale for these deformations will therefore be:

$$R_{min} \sim \frac{\gamma_t}{\sigma_{normal}(\rho_{AB})}.$$

This is the minimum length scale of deformation below which tension dominates and will damp out all active fluctuations. Deformations at or above this scale will have the opportunity to grow in amplitude and hence eventually lead to pinch-off of bits of target membrane.

The dependence of $\sigma_{normal}$ on the local antibody density is unknown, but one can make a reasonable assumption of its form. Below a critical density no engagement must occur between the macrophage and target, and hence no active stresses develop at the interface. Above a critical density we expect the active stresses to be non-zero and also scale with the antibody density and eventually saturate. Based on this, we assume the functional form:

$$\sigma_{normal}(\rho_{AB}) = \sigma_{max} \frac{\rho_{AB}}{\rho_{AB} + \rho_{bind}},$$

Where $\sigma_{max} \approx 150\ \mathrm{Pa}$, is the maximum scale of normal stresses measured in experiments[2], and $\rho_{bind}$ is the critical antibody density below which no interaction occurs between the macrophage and target. Based on our experiments we find that this density is ~100 μm[2].

One can similarly derive a second length scale which depends on the target membrane's resistance to local bending deformations due to active stresses, by balancing moments at the interface:

$$\sigma_{normal}(\rho_{AB})R_{bend}^2 \sim \frac{\kappa_t}{R_{bend}},$$

where $\kappa_t$ is the bending rigidity of the target membrane. This leads to the length scale of bend deformations:

$$R_{bend} \sim \left[\frac{\kappa_t}{\sigma_{normal}(\rho_{AB})}\right]^{\frac{1}{3}}.$$

This length scale can again be interpreted as a lower threshold below which membrane bending rigidity would preclude deformation. For typical values of bilayer bending stiffness $\kappa_t \approx 400\ k_B T$ and the scale of normal stresses measured at the macrophage-target interface $\sigma_{normal} \approx 100\ Pa$, this equation leads to a predicted deformation scale $\sim 0.1\ \mu\mathrm{m}$. For typical values of tension measured in experiments $\gamma_t \sim 10^{-2} - 10^{-1}$ mN/m, this means that the tension-mediated minimum length scale is greater and hence controls the scale of the deformations.

We use the above scaling laws to interpret our experimental data as follows. When $R_{min} \gg R$, where $R$ is the total interface size, the macrophage is unable to locally deform the target to pinch-off bites, and we predict that this biases the macrophage towards phagocytic behaviors. On the other hand, if $R_{min} \ll R$, there exists multiple length scales below the interface size that can be deformed and eventually pinched-off by the macrophage. We thus predict that when $R_{min} \ll R$, there is a bias towards trogocytic behaviors. Based on this one can derive a critical tension/antibody density that separates these distinct behaviors by setting $R_{min} \sim R$. Which gives:

$$\gamma_t|_{crit} \sim \sigma_{normal}(\rho_{AB})R.$$

For a typical macrophage-target interface scale of $\sim 1\ \mu m$ and $\sigma_{normal}(\rho_{AB}) \sim 50 - 150\ Pa$. This gives a critical target cortical tension that separates these behaviors of:

$\gamma_t|_{crit} \sim 0.1\ mN/m.$

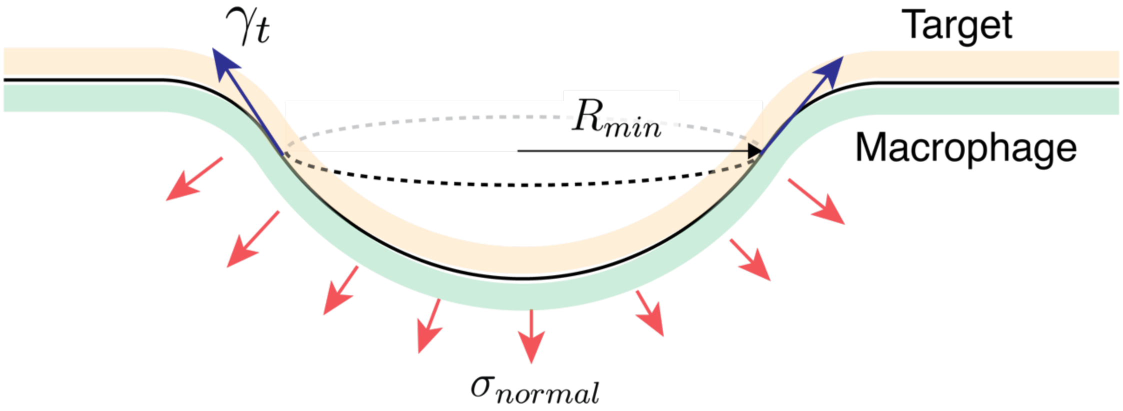


**Figure S8.** Balance of stresses at the phagocytic synapse between a macrophage and target cell.
